# Accurate and efficient protein sequence design through learning concise local environment of residues

**DOI:** 10.1101/2022.06.25.497605

**Authors:** Bin Huang, Tingwen Fan, Kaiyue Wang, Haicang Zhang, Chungong Yu, Shuyu Nie, Yangshuo Qi, Wei-Mou Zheng, Jian Han, Zheng Fan, Shiwei Sun, Sheng Ye, Huaiyi Yang, Dongbo Bu

## Abstract

Protein sequence design has been widely applied in rational protein engineering and increasing the design accuracy and efficiency is highly desired. Here we present ProDESIGN-LE, an accurate and efficient design approach, which adopts a concise but informative representation of residue’s local environment and trains a transformer to select an appropriate residue at a position from its local environment. ProDESIGN-LE iteratively applies the transformer on the positions in the target structure, eventually acquiring a designed sequence with all residues fitting well with their local environments. ProDESIGN-LE designed sequences for 68 naturally occurring and 129 hallucinated proteins within 20 seconds per protein on average, and the predicted structures from the designed sequences perfectly resemble the target structures with state-of-the-art average TM-score exceeding 0.80. We further experimentally validated ProDESIGN-LE by designing five sequences for an enzyme, chloramphenicol *O*-acetyltransferase type III (CAT III), and recombinantly expressing the proteins in *E. coli*. Of these proteins, three exhibited excellent solubility, and one yielded monomeric species with circular dichroism spectra consistent with the natural CAT III protein.

## Introduction

Protein sequence design, the inverse of protein folding, aims to design protein sequences that fold into a desired backbone structure (1). Computational approaches to protein sequence design have been widely used in rational protein engineering, including design of functional enzymes (2, 3), drugs (4, 5), and vaccines (6, 7). Despite the extraordinary advances (8–10), increasing the efficiency and accuracy of protein sequence design still remains a great challenge.

Protein sequence design can be accomplished using a concurrent strategy, in which the all residues of the protein are determined simultaneously. Most approaches of this type seek to exploit the inter-residue distances derived from the target structure for sequence design. For example, SPROF views the inter-residue distance map as an image with the designed sequence as its caption. In an analogy to the task of image captioning, SPROF applies a convolutional neural network (CNN) to infer protein sequence from the inter-residue distance map (11). ProteinSolver, another representative approach, uses a deep graph neural network to find a protein sequence that satisfies as many inter-residue constraints derived from the target structure as possible (12). These approaches are usually very efficient as they assign all residues of the protein simultaneously.

Protein sequence can also be designed in an iterative fashion — at each iteration step, the residue at a randomly selected position is mutated to improve the fitness between the entire protein sequence and the target structure. To measure the fitness, Rosetta sequence design’s fixed backbone (FixBB) protocol (13) uses Rosetta (14) energy, which is a human-crafted energy function consisting of dozens of energy terms, including physics-based terms such as van der Waals forces and solvation energy, and knowledge-based terms such as torsion angle preference. Notably, neural networks have been widely used by this strategy: Anand et al. proposed to learn potential directly from existing protein structures using a 3D-CNN model (9); SPIN2 (15), DenseCPD (16), and ProDCoNN (17) use deep neural networks to predict the most likely substituting residue type at a position conditioned by its surrounding structural features.

Due to the local nature of the residue-residue interactions within proteins, most of the existing approaches exploit local environment of a certain residue in a target structure (referred to as *target residue* hereinafter), including geometry features (e.g., relative position of surrounding residues) and chemical context (e.g., residue type of neighboring residues and solvent accessibility) to predicted the target residue. For example, DenseCPD and 3D-CNN draw a 20 × 20 × 20 Å^3^ box centered on a target residue as boundary of a local environment. DenseCPD uses four backbone atoms, i.e., *N,C,C_α_,O,* and *C_β_* in local environments to predict the most likely amino acid types for target residues, which are then fed into FixBB as additional design constraints, aiming to reduce sequence search space during design. In contrast, 3D-CNN uses all atoms within this box, including both backbone and rotamer atoms. It should be noted that when considering all atoms, we have to rebuild rotamers and evaluate the full-atom structures at each iteration step, which will greatly decrease the design efficiency.

Here, we present ProDESIGN-LE, an accurate and efficient approach to protein sequence design (Fig. 1A). The rational underlying our approach is that a designed protein, if every composing residue fits well with its local environment defined by the target structure and neighboring residues, is expected to fold into a structure globally resembling the target structure. ProDESIGN-LE uses a concise but informative representation of local environment, which describes the chemical context of a target residue using residue type rather than atoms of neighboring residues. Inspired by previous studies on protein structure prediction (18, 19), we uses the rotations from a target residue to its neighboring residues as a critical component of local environment’s representation (Fig. 1C and Supplementary Fig. 1). We further designed a transformer to learn the dependency of a residue on its local environment. ProDESIGN-LE iteratively applies the trained transformer to select an appropriate residue at a random position of the target structure, and updates the local environments of the neighboring residues accordingly, eventually acquiring a designed sequence with all residues fitting well with their own local environments. Our approach does not require frequent rebuilding the full-atom structures at the intermediate design steps, thus greatly improving design efficiency. In addition, by using the residue types to characterize the chemical context, our approach achieves higher accuracy compared with design approaches considering backbone atoms alone.

**Fig. 1.**
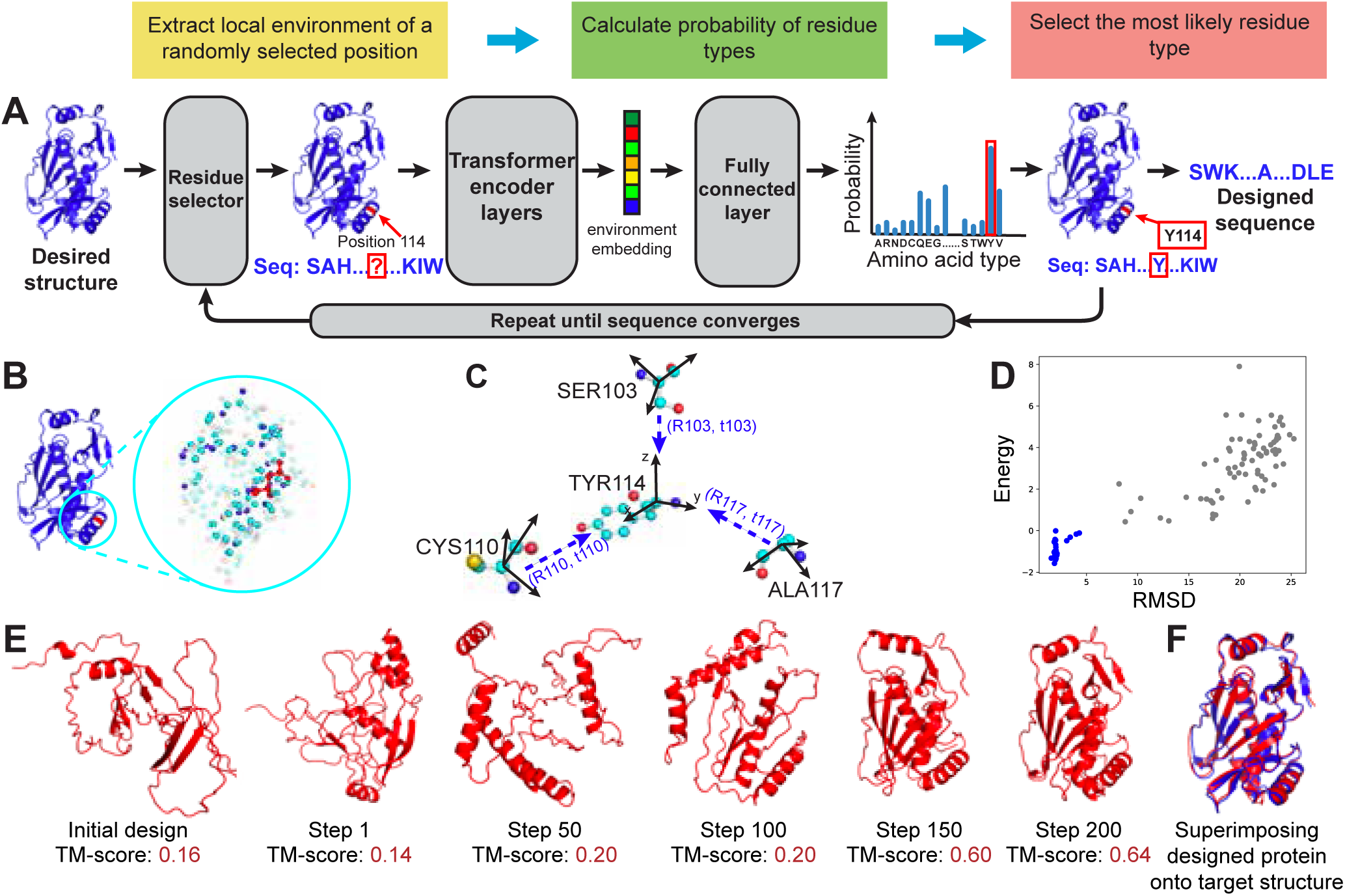
Overview of protein sequence design using ProDESIGN-LE. **A**, To design a sequence for a desired target backbone structure, ProDESIGN-LE starts from a random sequence and then iteratively selects the appropriate residue type for a random position, e.g., the local environment of position 114 is fed into transformer encoder and FC layer, yielding a distribution over 20 residue types. ProDEISGN-LE selects the most likely amino acid `TYR` and thus mutates the design sequence accordingly. ProDESIGN-LE repeats these steps until all residues fits well with their local environments, eventually acquiring a designed sequence. **B,** An example of the full-atom local environment around a target residue (in red), which contains all atoms within a sphere with a pre-defined radius centered at the residue. **C,** The concise but informative representation of local environment used by ProDESIGN-LE considers relative positions of neighboring residues. For the residue `TYR114`, its three neighbors `SER103, CYS110`, and `ALA117` are shown here. For each residue, we construct a local frame with 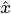 and *ŷ* being the result of applying the Gram–Schmidt process to 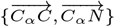, *ẑ* being 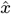 × *ŷ*. We then calculate a 3 × 3 transform matrix *R* and a 3-dim translation vector *t* for each neighbor with respect to `TYR114`. **D,** Energy vs. RMSD plot of the predicted structure for the intermediate sequences during redesigning protein CAT III. Here, RMSD measures the proximity of the predicted structure to the target structure. The structures with smaller RMSD usually have lower energy, especially for the native-like proteins with RMSD less than 5Å (blue dots). **E,** The design process for protein CAT III. The initial random sequence has its predicted structure deviating greatly from the target structure (TM-score: 0.16). After 200 rounds of iteration, ProDESIGN-LE acquires a design with associated structure perfectly matching the target structure (TM-score: 0.64). **F,** The superimposition of the predicted structure for the designed sequence (red) with the target structure (blue).

We assessed ProDESIGN-LE *in silico* using 68 naturally occurring proteins and 129 hallucinated proteins, and compare its performance with FixBB, 3D-CNN, and Protein-Solver. We further experimentally validated ProDESIGN-LE by designing five sequences for an enzyme, chloramphenicol *O*-acetyltransferase type III (CAT III) and recombinantly expressing the designed proteins in *E. coli*. The *in silico* assessing and experimental characterizing results clearly demonstrate the accuracy and efficiency of ProDESIGN-LE in protein sequence design.

## Results and discussion

### The concept of ProDESIGN-LE approach

We aim to design a sequence *S* that can fold into a desired backbone structure *B,* which is specified using coordinates of each residue’s three backbone atoms, *N,Cα* and *C*. Formally, the *i*th residue’s position is represented as *B_i_* = (*N_i_*, *Cα_i_*, *C_i_*), where *N_i_*,*Cα_i_*,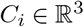 denote the 3D coordinates of these atoms, respectively. The design is denoted as *S* = *s*_1_*s*_2_… *s_n_*, where *s_i_* and *n* denote the *i*th residue and sequence length, respectively.

We represent the fitness of the designed sequence *S* with the desired backbone structure *B* as a conditional distribution *P*(*S*|*B*), which is approximated using the fitness of each residue with its local environment. The local environment of a residue describes its chemical context using the residue type of its neighbors and describes its geometric features using the relative positions of the neighbors. Formally, we define the *i*th residue’s local environment as *env_i_* = {(*s_j_*, *B_j_* ϴ *B_i_*, *j* – *i*)|||*Cα_i_* – *Cα_j_* || ≤ *T*}, where *T* represents a distance cut-off (set as 12 Å in the study), and the operator ϴ calculates the relative position of two residues, including relative rotation and translation. Using the relative position with respect to the target residue *B_j_* ϴ *B_i_* instead of the original coordinate *B_j_* gives our approach advantage of the rotation and translation invariance.

We further represent the fitness of a residue with its surrounding local environment as *P*(*s_i_*|*env_i_*). Using these notations, we approximate *P*(*S*|*B*) as its pseudo-likelihood:

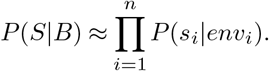

Our design algorithm aims to find a sequence *S* that maximizes the fitness *P*(*S*|*B*). We accomplish this objective through decomposing it into residue-wise sub-objectives, i.e., maximizing each residues’ fitness with its surrounding local environment *P*(*s_i_*|*env_i_*). In particular, our algorithm starts from a random sequence and then iteratively executes the following three steps for improvement: 1) randomly selects a position *i*, 2) calculates the conditional probability *P*(*a*|*env_i_*) for each possible residue type *a,* and mutates *s_i_* to be the most likely residue type, i.e., *s_i_ ← argmax_a_P*(*a*|*env_i_*), and 3) all neighbors of the *i*-th residue have their local environments automatically updated after mutating *s_i_*. These three steps are iterated until the fitness *P*(*S*|*B*) converges.

In our approach, we learn the conditional distribution *P*(*s_i_* |*env_i_*) using a transformer as classifier, which is trained on a subset of PDB40 dataset (see Methods for details). The geometrical features in local environment *env_i_*, i.e., the relative position of two residues *B_j_* ϴ *B_i_*, is represented as a 3 × 3 rotation matrix together with a 3 × 1 translation vector. We flat the rotation matrix into a 9 × 1 vector, catenate it with other features (e.g., one-hot encoding of residue type and sequence position) as input to the classifier (for details see Supplementary Fig. 1).

We evaluated ProDESIGN-LE using two ways, i.e., comparing the predicted structures of the designed sequences with target structures, and recombinantly expressing the designed proteins in *E. coli* and experimentally characterizing them. In the study, we predict structure for the designed sequence using AlphaFold2 (19) and our inhouse software ProFOLD Zero, an improved version of ProFOLD (20). ProFOLD Zero is suitable for the task of sequence design as it was specially designed for the structure prediction of single protein sequence without requirement of multiple sequence alignment.

Using CAT III protein as an example, we describe the major steps and operations of ProDESIGN-LE as below (Fig. 1E). We aim to design a protein with the backbone structure of the CAT III from *E. coli* as the target (PDB entry: 6X7Q, 212 a.a.). Initially, ProDESIGN-LE set the protein sequence randomly as `SAHIP....QKIW` (the entire sequences of the initial design and intermediate designs are provided in Supplementary Text 1). As expected, the 3D structure predicted from this random sequence deviates significantly from the target structure (TM-score: 0.16). At the first step, ProDESIGN-LE selected position 114, whose local environment involves the information of its three neighbors, i.e., `SER103, CYS110`, and `ALA117`. By feeding this local environment into the geometry transformer and the follow-up fully connected layer, ProDESIGN-LE calculated the probability of all residue types and selected the most likely residue type (`TYR`: 0.30) to replace the original residue type at this position. After repeating this procedure 200 times, ProDESIGN-LE yielded a designed sequence SWRTVD….SDPE with its predicted structure in perfect agreement with the target structure (TM-score: 0.64). In contrast, the predicted structure of the native sequence achieves a TM-score of 0.66. Fig. 1D plots the energy of the predicted structures versus their RMSDs with respect to the target structure, which suggests that the structures with smaller RMSD usually have lower energy, especially for the native-like proteins with RMSD less than 5 Å (blue dots).

### Redesign naturally occurring proteins using ProDE-SIGN-LE

Using 68 naturally occurring protein domains extracted from the CASP14 dataset as representatives (see Supplementary Text 2 for full list), we evaluated ProDESIGN-LE and compared it with the widely-used design approaches, including 3D-CNN model, ProteinSolver, and FixBB.

The evaluation criteria include: (1) *sequence matching* (aka sequence identity): the sequence identity between the designed sequence and native sequence of the target structure, (2) *structure matching:* the structure similarity (measured using TM-score) between the target structure and the predicted structure of the designed sequence. (3) *sequencestructure matching:* we further evaluated the fitness between the designed sequence and the target structure through building and assessing a threading structure. The threading structure was built as performed by template-based modelling (21), i.e., complementing the target backbone structures with the sidechains determined by designed sequences and then fine-tuning the sidechain conformation using Rosetta relax protocol (22). The energy of the resultant threading structure is used as measurement of the fitness between the designed sequence and the target backbone structure, and an ideal designed sequence is expected to have a low energy.

We show the design for protein T1093-D3 as a concrete example (Supplementary Fig. 2). Protein T1093-D3 contains a total of 106 residue with 3 *α*-helices and 10 *β*-strands in its native structure. The sequence identity between the design and protein T1093-D3 is 0.28. For 92 out of the 106 residues, the predicted structure is in perfect agreement with the target structure (mean *C_α_* RMSD: 0.84 Å). The designs for 20 proteins are shown in Supplementary Figure 3 as representatives.

For the 68 CASP14 proteins, the sequence designs by ProDESIGN-LE achieved an mean sequence identify of 0.33, at the same level with ProteinSolver (0.32), 3D-CNN (0.28), and FixBB (0.28). In contrast, the predicted structures of these designs achieved a mean TM-score of 0.84, which is significantly close to that of the predicted structure using the native sequence (0.88) and higher than all other design approaches including ProteinSolver (0.75), 3D-CNN (0.74), and FixBB (0.64). In addition, the threading structures generated using the designs by ProDESIGN-LE have lower energy than other design approaches (Fig. 2A-C).

**Fig. 2.**
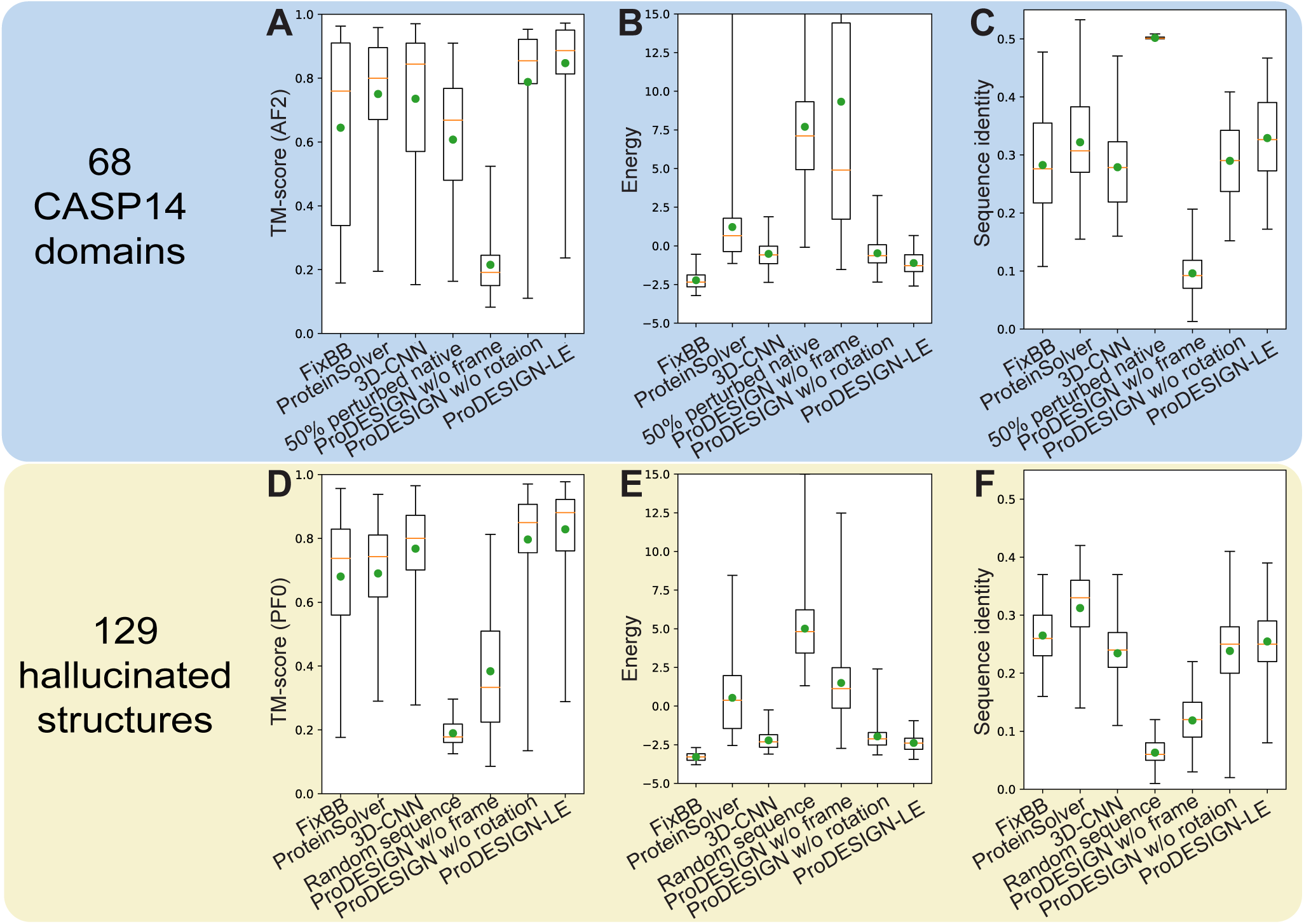
Assessing *in silico* the designed sequences for 68 naturally occurring (A, B, C) and 129 hallucinated proteins. **(D, E, F)**. We designed sequences for these proteins using FixBB, ProteinSolver, 3D-CNN, and ProDESIGN-LE and assessed the designed sequences using three metrics: (1) the sequence identity between the designed sequence and native sequence of the target structure (**C, F**), (2) the structure similarity (measured using TM-score) between the target structure and the predicted structure of the designed sequence (**A, D**), (3) we further built a threading structure through complementing the target backbone structure with the sidechains determined by designed sequences. The energy of the resultant threading structure is used as measurement of the fitness between the designed sequence and the target backbone structure (**B, E**). We used AlphaFold2 (denoted as AF2) to predict structures for the naturally occurring proteins and used ProFOLD Zero (denoted as PF0) to predict structures for the hallucinated proteins.

We also carried out ablation analysis of ProDESIGN-LE through evaluating the variants with coordinate frames or rotations of neighboring residues removed from local environments. As shown in Fig. 2 A and D, the coordinate frames of neighboring residues are critical information of local environments.

### Assessing the generalization of ProDESIGN-LE to hallucinated proteins

We further assessed the generalization of ProDESIGN-LE using 129 hallucinated proteins, which were created through inverting the neural networks trained to predict structures from sequences (23). These hallucinated proteins serve as ideal test data for evaluating the generalization of protein design approaches as they are de *novo* and unrelated to the proteins used to train the neural networks.

For the 129 hallucinated proteins, all approaches yielded designs with sequence identity at the same level. However, the structures predicted using ProFOLD Zero from the designs by ProDESIGN-LE achieved an average TM-score of 0.83, which is considerably better than the designs by ProteinSolver (0.69), 3D-CNN (0.77), and FixBB (0.68) (Fig. 2D, F). We repeated this experiment using AlphaFold2 as prediction tool and achieved similar observation (Supplementary Fig. 4).

In addition, when using the design by ProDESIGN-LE, the average energy of the resultant threading structures is −238.57, which is close to those generated using FixBB (−328.72) and lower than those generated using ProteinSolver (53.25), 3D-CNN (−221.50), and random sequences (500.88). Thus, compared with other approaches, the designs by FixBB and ProDESIGN-LE are more compatible with the hallucinated structures (Fig. 2E).

ProDESIGN-LE also showed considerably high time efficiency: ProDESIGN-LE accomplishes sequence design for CAT III with 212 residues within 25 s on an ordinary GPU (Nvidia RTX 3090, memory: 24GB), which is greatly faster than FixBB (348 s). For the 68 naturally occurring and 129 hallucinated proteins, ProDESIGN-LE accomplished sequence design within 20 seconds per protein on average. The running time of ProDESIGN-LE increases quadratically with the length of target structure as expected (Supplementary Fig. 5).

Together, these results demonstrate that ProDESIGN-LE can be used to efficiently design both naturally occurring proteins and hallucinated proteins.

### Assessing the effectiveness of local environment on predicting residue type

An ideal definition of local environment around a residue should effectively describe the preference of residue types in the environment. To assess the effectiveness of the local environment used by ProDESIGN-LE, we calculated accuracy of the predicted residue types in each local environment. We also investigated the correlation between prediction accuracy and entropy of the predicted distribution of residue types. We used the negative entropy as confidence score of the prediction.

We sorted all 758,160 local environments extracted from the test set according to their confidence scores, and then calculated the accuracy of the predicted residue types that might appear in these local environments. The top 10% of local environments with the highest confidence score achieved a top-1 and top-5 prediction accuracy of 0.902 and 0.982, respectively, i.e., for 90.2% and 98.2% of these local environments, ProDESIGN-LE can successfully rank the groundtruth residue type as top 1 and top 5, respectively (Fig. 3A). Even for the top 50% of local environments with high confidence score, the prediction accuracy is still considerably high (top-1 accuracy: 0.567, top-5 accuracy: 0.892).

**Fig. 3.**
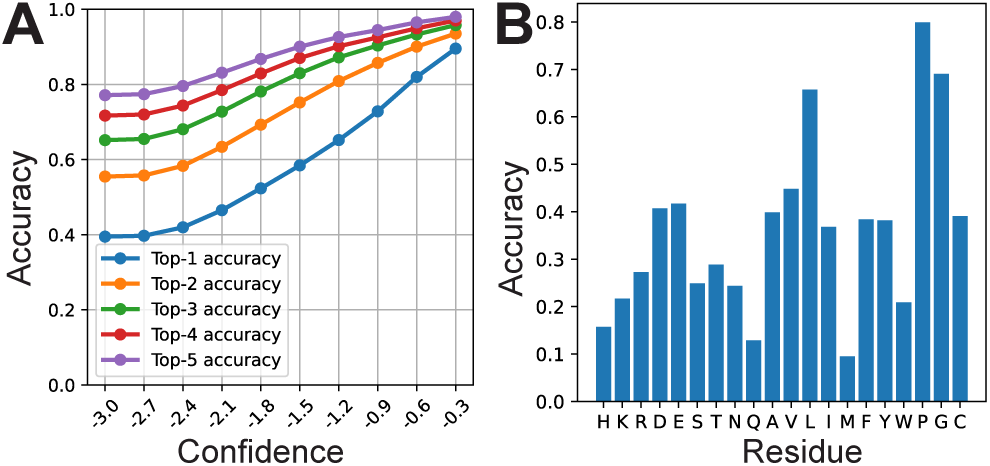
Accuracy of prediction of a residue according its local environment. **A,** ProDESIGN-LE predicts a distribution over 20 residue types for a target residue, and assigns each residue type with a confidence score. We calculate the top-K (*K* =1, 2, 3, 4 and 5) accuracy of the predicted residue types exceeding a confidence score cut-off (*x* axis). **B,** The relationship between the prediction accuracy and the ground-truth residue type extracted from the native sequence of the protein. **C,** The confusion matrix shows the ground-truth residue type and the predicted distribution over all possible 20 residue types.

In-depth examination suggested that the prediction ability of a local environment is tightly correlated with the residue it surrounds. As shown in Supplementary Fig. 6, the prediction confidence score of local environments exhibits a substantial correlation with the solvent accessibility of central residues (Pearson correlation coefficient: −0.68). Moreover, the prediction accuracy is high in the case that the groundtruth residue is `Pro` (0.80) or `Gly` (0.69), implying that the local environment around these residues are more constrained (Fig. 3B). This correspond well with previous studies which showed proline and glycine confer unique structural constraints on backbone due to proline’s distinctive cyclic structure and glycine’s lack of side chain (24, 25), thus result in the distinctive local environments around them. In contrast, the prediction accuracy is relatively low in the case that the ground-truth residue is `Met` (0.09): methionine is frequently predicted as leucine (Supplementary Fig. 7), which coincides with its chemical property.

It should be noted that the prediction accuracy increases monotonically with confidence score (Fig. 3A), thus enabling the use of confidence score as an effective index of the reliability of prediction.

### Experimental characterization of designed sequences of CAT III

CAT III is an enzyme that confers resistance of antibiotic chloramphenicol to certain bacterium (26). The functional form of CAT III is a homotrimer, of which the substrate pockets lie between two adjacent protomers. We chose the CAT III from *E. coli* as the design target, which consists of 212 residues, forming five *α*-helices, two 3_10_-helices, and two *β*-sheets (Fig. 1F).

We used CAT III as a representative example to evaluate ProDESIGN-LE through experimentally characterizing its designs for this protein. In particular, we executed ProDESIGN-LE using the backbone structure of CAT III as a target structure, and acquired 5 designed sequences (denoted as CAT-h1, CAT-h2, CAT-h3, CAT-h4, CAT-h5) as results. To experimentally characterize the designed sequences, we synthesized their coding genes, and recombinantly expressed the corresponding proteins in *E. coli*. The recombinant proteins were separated using SDS-PAGE. As shown in Supplementary Fig. 8, all 5 proteins were successfully expressed, which matches well with mass of the 6xHis-tagged protein at around 29 kDa. In addition, three out of the five proteins, CAT-h2, CAT-h3 and CAT-h4, were expressed as soluble form.

We further purified the soluble designs by Ni-affinity chromatography for in-depth analysis. Sedimentation velocity experiment of the 6xHis-tagged CAT-h2 yielded a peak at 28.9 kDa, indicating that CAT-h2 exists in a monomeric form with a mass of approximately 25 kDa (Supplementary Fig. 9). The designed CAT-h2 was highly thermostable with a melting temperature of 72.5 °C, comparable with CAT III (74.8 °C, Fig. 4A and Supplementary Table 1). In addition, structural superimposition of the predicted structure of CAT-h2 and the target structure revealed a 4.02 Å *C_α_* RMSD over 212 residues. Here, we examined design CAT-h2 by circular dichroism (CD) spectroscopy. The CD spectra of CAT-h2 were in good agreement with those acquired from CAT III (Fig. 4B), showing characteristic profiles of *α/β* proteins (Supplementary Tables 2, 3).

**Fig. 4.**
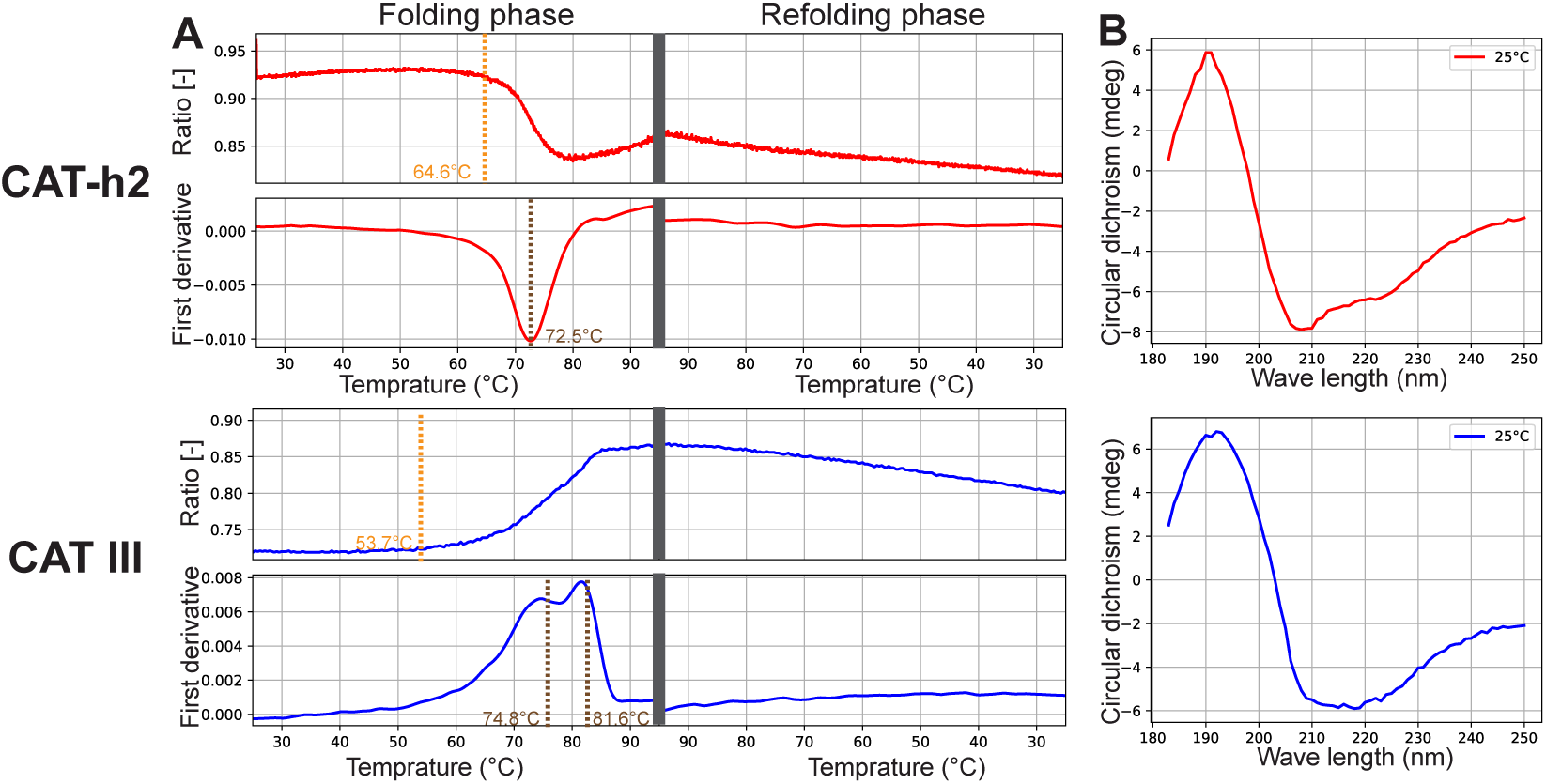
Experimental characterization of the designed protein (CAT-h2) and natural protein (CAT III). **Column A,** Thermostable analysis of the two proteins by nanoDSF measurement. The designed protein CAT-h2: the onset denaturation temperature of protein is 64.6 °C and the folding Tm value is 72.5 °C; The natural protein CAT III: the onset denaturation temperature of protein is 53.7 °C and the folding Tm values is 74.8 °C. Ratio: 350nm/330nm fluorescence intensity. **Column B,** Circular dichroism spectra of the proteins from 185 to 260nm at 25 °C. The designed protein CAT-h2 (red) exhibited circular dichroism spectra consistent with the natural protein CAT III (blue).

Together, these results clearly demonstrate that the designed proteins by ProDESIGN-LE can be successfully expressed and can fold into stable structures with the desired secondary structures.

## Conclusion

The *in silico* assessments and experimental characterization results presented here for protein sequence design by ProDESIGN-LE have highlighted the special features of learning the concise but effective representation of local environments around residues. The results have also highlighted the superiority of the design paradigm used by ProDESIGN-LE, in which a protein with expected global structure is designed through iteratively selecting an appropriate residue at a random position to fit well with its local environment. The accuracy and efficiency of ProDESIGN-LE have been clearly demonstrated using both naturally occurring and hallucinated proteins as representatives, along with experimental characterization of the designed proteins expressed in *E. coli*.

The present ProDESIGN-LE considers the proximity between the structure of the designs with the target structures only but pays no attention to the biochemical characteristics of the designed proteins. At present, of the five designed proteins for CAT III, three showed excellent solubility but two were insoluable. One possible reason of the two insoluable proteins might be the overuse of hydrophobic residues on protein surface (Supplementary Fig. 10). Thus, a possible improvement might be posing restrictions on the use of residues in the surface and core. In addition, determining how to design multimer with significant affinity among subunits is also highly desired. This might be accomplished through adding a neural network module to predict affinity and adding it into the objective function. In future studies, these will be incorporated into ProDESIGN-LE. The algorithm and operations of ProDESIGN-LE can be readily extended to further improve success rate of design without significant modifications of the basic ideas.

We anticipate that our work on protein sequence design by ProDESIGN-LE with improved accuracy and efficiency will facilitate the rational protein engineering.

## Methods

### Network architecture

ProDESIGN-LE consists of a 3-layer transformer encoder that takes local environments as input and yields their embeddings as output (27). ProDESIGN-LE further uses a fully-connected layer to transform the embeddings into a distribution over 20 residue types. We implemented ProDESING-LE in Python (ver. 3.8.8) and PyTorch (ver. 1.9.1).

### Model training

ProDESIGN-LE uses the cross-entropy loss function as its optimization objective function. We train our model with a batch size of 1000 using Adam optimizer (*β*_1_ = 0.9, *β*_2_ = 0.999) with learning rate 1 × 10^-3^ (28). The training process costs about 2 hours on a single Nvidia RTX 3090.

### Datasets

We train and test the transformer using the structures collected in the PDB40 database (29). We excluded the structures that contain any DNA/RNA chain or structures contain multiple models. To avoid potential data leaking, we constructed test set with structures that have no similar structures in training set, this is achieved via MMseq2 by filtering structures in test set with sequences similar (e-value < 1 × 10^-3^) to any sequence in training set.

Each sample in training and test set represents a residue of the protein structures obtained as mentioned above. For each residue, we describe its local environment as the following features: 1) the residue type of neighboring residues in one-hot encoding, 2) the relative 3D position of each neighboring residue with respect to the target residue, represented as a rotation matrix and a translation vector, 3) the distance between a neighbor residue with the target residue in sequence.

As results, we acquired a training set containing 5,867,488 residues extracted from 9,995 protein structures and a test set containing 758,160 residues extracted from 401 proteins structures.

We test the performance of ProDESING-LE in sequence design using 68 naturally occurring proteins extracted from CASP14 and 129 hallucinated structures (23).

### Predicting 3D structure for the designed sequences

We evaluate a designed sequence using the proximity between the predicted structure of the designed sequence and the target structure. Here, we predict 3D structures for the designed sequences through running AlphaFold2 with its default setting on a template released before 2020-02-25.

We further apply ProFOLD Zero, an improved version of ProFOLD (20), to predict structures for the hallucinated proteins. ProFOLD Zero is suitable for the task of sequence design as it was specially designed for the structure prediction of single protein sequence without requirement of multiple sequence alignment.

### Protein expression and purification

Coding DNA sequences of designed proteins were constructed into the *BamH I* and *Xho I* sites of pET-28a(+). *E. coli* BL21 (DE3) cells were transformed with the plasmids. Protein expression was induced at an *OD_600_* between 0.6 and 0.8 with 210 μmol/L IPTG for 16 h at 16 °C. Cells were harvested and sonicated in a lysis buffer containing 100 mmol/L Tris-HCl and 100 mmol/L NaCl at pH 8.5. The soluble supernatant was purified by Ni-affinity chromatography and collected in pH 8.5 50 mmol/L imidazole buffer. The purified protein was sealed in a dialysis bag at 4 °C, replaced with 400 mL 2 mmol/L Hepes buffer for 3 h, and dialyzed 3 times to avoid Tris and NaCl in the sample.

### Analyzing protein thermal stability

The protein thermal stability was analyzed via NanodsF-Prometheus NT.48 device by following the method used by Ref. Magnusson et al. 30. The protein samples were prepared in buffer containing 100 mmol/L NaCl, 10 mmol/L Tris, pH 8.5 with a protein concentration of 0.5 mg/mL. The 10 μL CFEs of samples was loaded in capillaries; the temperature gradient of 1 °C/min from 25 to 95 °C was applied and the intrinsic protein fluorescence at 330 and 350 nm were recorded. The nanoDSC scans were background-corrected and analyzed with Launch NanoAnalyze software.

### Circular dichroism spectroscopy

The circular dichroism spectra of the designed CAT proteins and natural CAT protein (PDB entry: 6X7Q) were determined by a Chirascan V100 Circular dichroism spectrometer (Applied Photophysics, https://www.photophysics.com/Britain) (31). The path length and volume of Quartz cells were 1 mm and 200 μL, respectively. The time-per-point was 0.5 s, the scanning step was 1 nm, and the scanning ranged from 185 to 280 nm. The spectrum was calibrated with Hepes buffer (2 mmol/L, pH 8.5). The ultraviolet absorbance of CAT protein was measured using a circular dichroism spectrometer, and the absorbance was in the range of 0.6 to 1.2, indicating that the signal-to-noise ratio was suitable for analysis. The 100 μL samples were typically loaded at concentrations of 0.15 to 0.2 mg/mL in pH 8.5 2 mmol/L Hepes buffer into a quartz cell. The results were taken as CD ellipticity in mdeg. The percentages of *α*-helix, *β*-sheets, and random coils designed CAT proteins and control CAT proteins were calculated using the CDNN soft and net using 23 basespectra (advanced CD spectra).

### Analytical ultracentrifugation

Sedimentation velocity experiments were performed in a ProteomeLab XL-I analytical ultracentrifuge (Beckman Coulter, Brea, CA)(32), equipped with AN-60Ti rotor (4-holes) and conventional double-sector aluminum centerpieces of 12mm optical path length, loaded with 380 μL sample and 400 μL buffer (10 mmol/L Tris-HCl, pH 7.0, 100 mmol/L NaCl). Before the run, the rotor was equilibrated for approximately 1 h at 20 ^°^C in the centrifuge. Then experiments were carried out at 20 ^°^C and 41000 rpm, using continuous scan mode and radial spacing of 0.003 cm. Scans were collected in 3 min intervals at 280nm. The fitting of absorbance versus cell radius data was performed using SEDFIT software (https://sedfitsedphat.nibib.nih.gov/software/default.aspx) and continuous sedimentation coefficient distribution c(s) model, covering range of 0 to 20 S. Biophysical parameters of the buffer are: density *ρ* = 1.006 g/cm^3^, viscosity *η* = 0.01031P, and the parameters of proteins are: partial specific volume V-bar = 0.73000 cm^3^/g.

### Statistical analysis

All experiments were independently carried out at least three times and the results were expressed as mean ± standard deviation (SD).

## Acknowledgements

We would like to thank Haipeng Gong and Luhua Lai for invaluable suggestions and comments. We would also like to thank the National Key Research and Development Program of China (under grant 2020YFA0907000) and the National Natural Science Foundation of China (under grants 31871287, 62072435) for providing financial supports to the study.

