## Supplementary File 1 for "Accurate and efficient protein sequence design through learning concise local environment of residues"

### Contents


**Supplementary Text 1. The major steps of the design of CAT III** 3

**Supplementary Text 2. The list of the 68 CASP14 domains used to assess ProDESIGN-LE** 3

**Supplementary Table 1. Unfolding temperature Tm of the natural CAT III and the designed CAT-h3 proteins determined by by nanoDSF measurement** 4

**Supplementary Table 2. Secondary structure of the natural CAT III protein determined using circular dichroism spectrometry** 4

**Supplementary Table 3. Secondary structure of the designed CAT-h2 protein determined using circular dichroism spectrometry** 4

**Supplementary Figure 1: Architecture of the neural networks used by ProDESIGN-LE** 5

**Supplementary Figure 2. The predicted structure of the designed sequence by ProDESIGN-LE for protein T1093-D3** 6

**Supplementary Figure 3. Superimposition of the predicted structure of designed sequences onto the corresponding target structures** 7

**Supplementary Figure 4. *In silico* assessing the designed sequences for 129 hallucinated proteins** 8

**Supplementary Figure 5. Running time of ProDESIGN-LE for protein sequence design** 9

**Supplementary Figure 6. The relationship between the accuracy of the prediction of amino acid type and solvent accessibility of the target residue** 10

**Supplementary Figure 7. Confusion matrix of the transformer.** 11

**Supplementary Figure 8. SDS-PAGE analysis of the five designed CAT proteins crude extracts** 12

**Supplementary Figure 9. Sedimentation velocity analysis(SVA) of the designed CAT-h2 protein and the control CAT III protein** 13

**Supplementary Figure 10. The ratios of hydrophobic, hydrophilic and neutral residues in the core or on the surface of the designed proteins for CAT III** 14

### Supplementary Text 1. The major steps of the design of CAT III

Initial design (random sequence, 212a.a.):
SAHIPATFHQCASIIPNTWGHSMTDQVKGGKWIKRLSYSFSMCDRFMNDLWDEKPPQWGFTADWNRGSFAKCSHNSMIGDQILFTK
AEKSHYWLILPPICVSSMWPLSVCVQDKKMAAFTLIDYWYSNTNAVEECVCYMTNFSHVQADSGFIDKKIMCRADKKLIELTARWN
MSMESYWVDCHAEFGYICPIRKAYGQCYISYARLWSQKIW
Step 1:
SAHIPATFHQCASIIPNTWGHSMTDQVKGGKWIKRLSYSFSMCDRFMNDLWDEKPPQWGFTADWNRGSFAKCSHNSMIGDQILFTK
AEKSHYWLILPPICVSSMWPLSVCVQDYKMAAFTLIDYWYSNTNAVEECVCYMTNFSHVQADSGFIDKKIMCRADKKLIELTARWN
MSMESYWVDCHAEFGYICPIRKAYGQCYISYARLWSQKIW
Step 50:
SFRTPATFHQCASIIPNTWGHSMTDQVAGGKWLNRLDYSFSLCDRFMNDLWDEKWPQTGFINDINRGSFAVCNDNSMFGDQILFTK
AIYSHKLDVTPPISVSSSWPLDVCVQDYKMAADTYKDNPYSNPQAVEECVCYMTNFSHVQADSGFIDKKPMLRADKHPIELTARPN
MSMEDYWVDCHAQFNYICPIREDYDQYYISYQKLWSQKPW
Step 100:
SWRTPALFLQCDRIIPNTWGHSMPGQVAGTRWLNRLDYYFSLCRRDMNDLFDEVWPQTGFINDNDNFSFAVRNDQLMFFDQILPTV
TVYNPKLDVTPPISWKSSWDLDVCVQDYKMAADIYKDNPYKNPQGVPECVCYMTNFRHDEADSGFIDKKPMLRAQKHPIELTARPN
MSMEDLWVDCHIQFNYECPTQEDVDQWWINYQKLWSSKPE
Step 150:
SWRTVALFLSPDRERYNTWRNSMPGQAAGTRWLNRLDLYFSLSRRDMNLLFYEVYRQTGFINDNDKFSLTVRNDQLMFWDQVLPMV
TIYNPKLDVTPPIQWKSSWDIDEFVRDYKMAADIYKDNPLKVPQGVPEPVCYMTNFVHDRAYSGFEDKKPMLENQKHPIITTARPN
MRGEDLLLDCHIQFNYEVVTQEDVDTYWINYQKLLSSKPE
Step 200:
SWRTVDLFLSPERERYYYYRNIMPGQAAETRWLNRTDLYFSLSRSDMNLLDYEVWRQTLVINDNDKFSLRVRDDQLMSWDRVLPMV
TIRIPKTNTTPPLQWKFSWDINEFVRDYEMALKIYKDNPLKVPQGEPSEVRVMTNFVPDRYYSGFEDKKPNLENQKHPIITYARPN
RVGEDLLLPVSIQFNYAVVTKEDVDTLWINYQKLLLSDPE

### Supplementary Text 2. The list of the 68 CASP14 domains used to assess ProDESIGN-LE

We assessed ProDESIGN-LE using 68 naturally occurring proteins extracted from the CASP14 dataset shown below.

T1024-D1, T1025-D1, T1026-D1, T1028-D1, T1029-D1,
T1030-D1, T1030-D2, T1031-D1, T1032-D1, T1033-D1,
T1034-D1, T1035-D1, T1037-D1, T1038-D1, T1038-D2,
T1039-D1, T1040-D1, T1043-D1, T1045s1-D1, T1045s2-D1,
T1046s1-D1, T1046s2-D1, T1047s1-D1, T1049-D1, T1050-D1,
T1050-D2, T1050-D3, T1052-D1, T1052-D3, T1053-D2,
T1054-D1, T1055-D1, T1056-D1, T1060s2-D1, T1061-D0,
T1061-D2, T1061-D3, T1065s1-D1, T1065s2-D1, T1067-D1,
T1070-D1, T1070-D2, T1070-D3, T1070-D4, T1073-D1,
T1074-D1, T1078-D1, T1079-D1, T1080-D1, T1082-D1,
T1083-D1, T1084-D1, T1087-D1, T1089-D1, T1091-D1,
T1091-D2, T1091-D3, T1091-D4, T1092-D1, T1092-D2,
T1093-D1, T1093-D2, T1093-D3, T1094-D2, T1095-D1,
T1096-D1, T1096-D2, T1099-D1

### Supplementary Table 1. Unfolding temperature Tm of the natural CAT III and the designed CAT-h3 proteins determined by by nanoDSF measurement

| Sample | Onset 1 for Ratio (unfolding) | Inflection Point 1 for Ratio (Unfolding) |
|---|---|---|
| CAT III | $53.7 \pm 3.0\,°C$ | $74.8 \pm 0.1\,°C$/$81.6 \pm 0.5\,°C$ |
| CAT-h2 | $64.6 \pm 0.2\,°C$ | $72.5 \pm 0.1\,°C$ |

### Supplementary Table 2. Secondary structure of the natural CAT III protein determined using circular dichroism spectrometry

| | 180-260 nm | 185-260 nm | 190-260 nm | 195-260 nm | 200-260 nm | 205-260 nm | 210-260nm |
|---|---|---|---|---|---|---|---|
| $\alpha$-helix | 28.6% | 30.6% | 29.4% | 28.4% | 29.0% | 28.2% | 26.0% |
| Anti-parallel $\beta$-sheet | 12.0% | 11.0% | 12.3% | 14.8% | 12.6% | 13.2% | 14.7% |
| Parallel $\beta$-sheet | 6.6% | 7.2% | 7.3% | 6.6% | 6.2% | 5.9% | 5.6% |
| $\beta$-Turn | 16.2% | 15.4% | 16.3% | 16.6% | 16.3% | 16.5% | 17.2% |
| Coil | 35.8% | 37.9% | 35.4% | 34.1% | 34.0% | 34.8% | 32.9% |
| Total | 99% | 102.0% | 100.80% | 100.5% | 98.1% | 98.5% | 96.5% |

### Supplementary Table 3. Ssecondary structure of the designed CAT-h2 protein determined using circular dichroism spectrometry

| | 180-260 nm | 185-260 nm | 190-260 nm | 195-260 nm | 200-260 nm | 205-260 nm | 210-260 nm |
|---|---|---|---|---|---|---|---|
| $\alpha$-helix | 19.8% | 18.6% | 18.7% | 18.5% | 18.7% | 18.9% | 19.0% |
| Anti-parallel $\beta$-sheet | 21.7% | 23.7% | 24.1% | 24.4% | 24.3% | 24.3% | 24.0% |
| Parallel $\beta$-sheet | 5.7% | 5.6% | 5.5% | 5.7% | 5.60% | 5.6% | 5.5% |
| $\beta$-turn | 18.3% | 18.8% | 18.9% | 19.0% | 19.1% | 19.1% | 19.0% |
| Coil | 34.9% | 34.1% | 33.8% | 34.3% | 34.7% | 34.2% | 34.3% |
| Total | 100.3% | 100.8% | 101.0% | 101.9% | 102.4% | 102.1% | 101.9% |

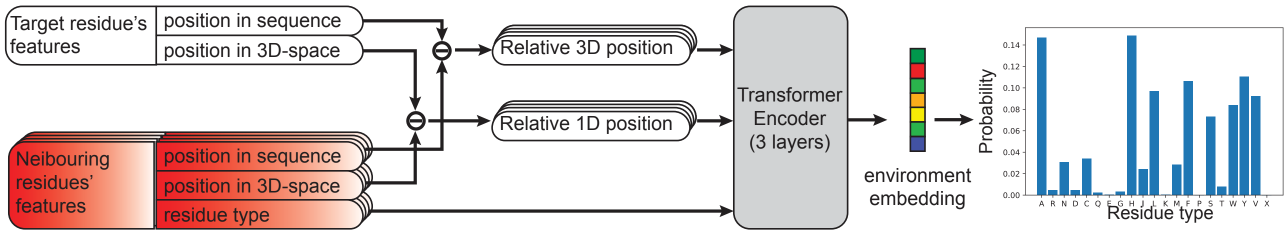


**Supplementary Figure 1. Architecture of the neural networks used by ProDESIGN-LE**

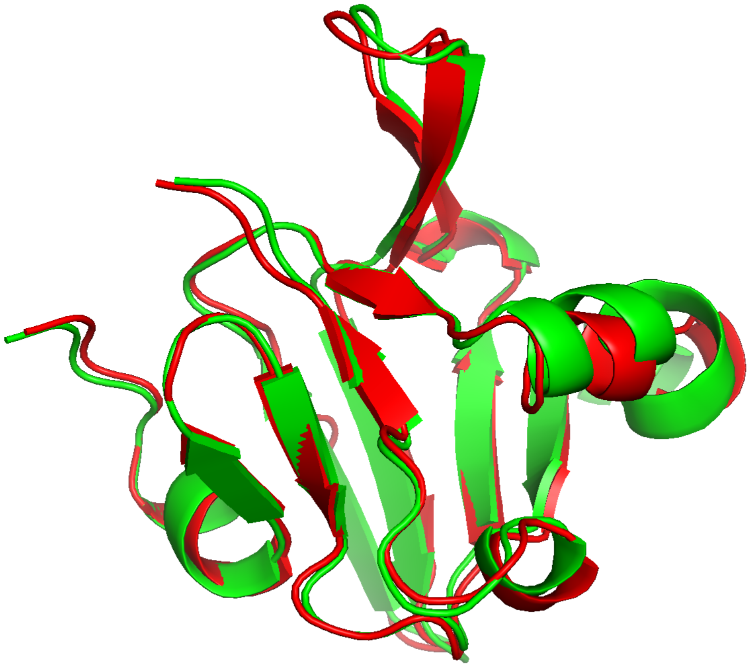

**Supplementary Figure 2: The predicted structure of the designed sequence by ProDESIGN-LE for protein T1093-D3**

The TM-score between the native structure of protein T1093-D3 (red) and the predicted structure (green) of the designed sequence by ProDESIGN-LE is 0.89.

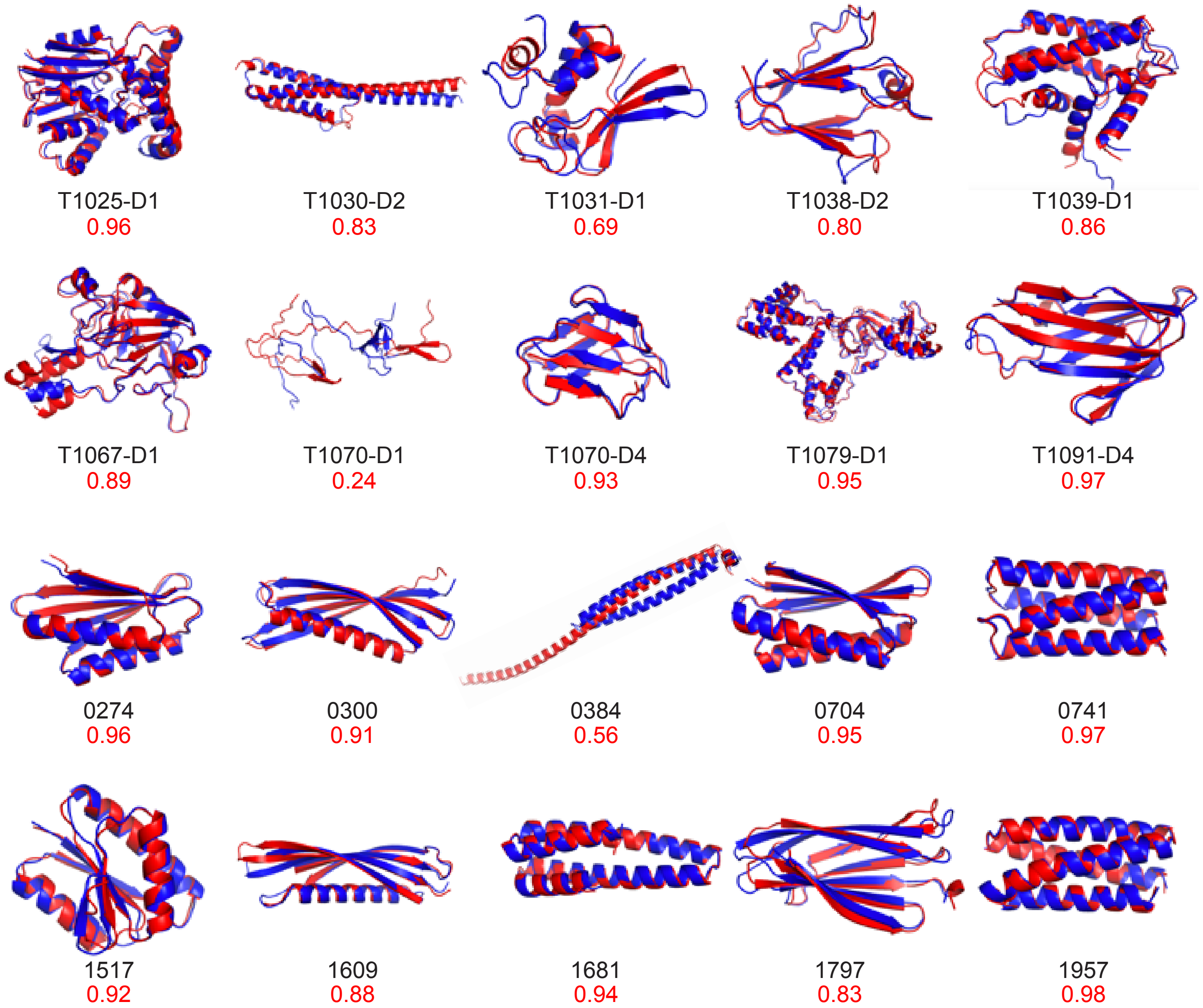


**Supplementary Figure 3. Superimposition of the predicted structures of designed sequences onto corresponding target structures**

Ten CASP14 proteins and ten hallucinated proteins are shown here as representatives. The TM-score between the predicted structure (red) and the corresponding target structure (blue) is shown below the superimposition diagram.

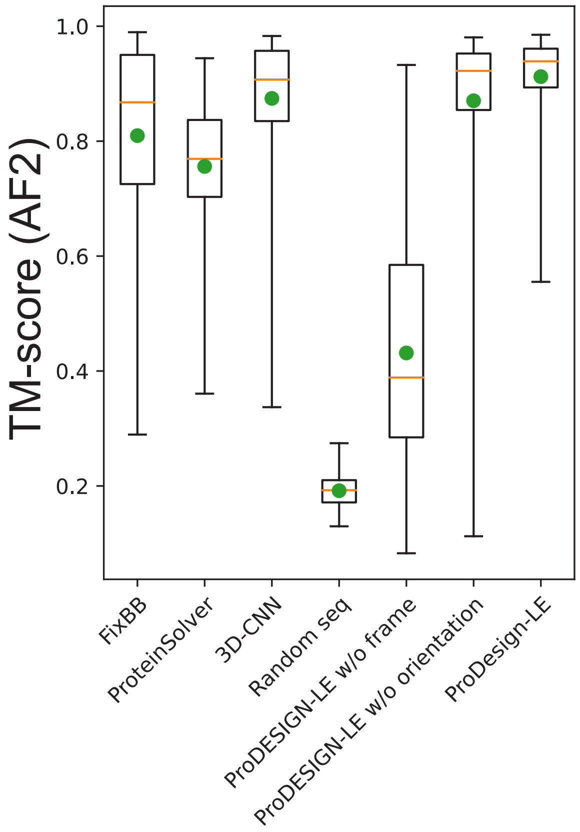


**Supplementary Figure 4. *In silico* assessing the designed sequences for 129 hallucinated proteins**

Using AlpahFold2, we predicted structures for the designed sequences of 129 *de novo* hallucinated structures by Rosetta-Fixbb, ProteinSolver, 3D-CNN, and ProDEDIGN-LE. We also predicted the structures using ProFOLD Zero (see Fig. 2D for details).

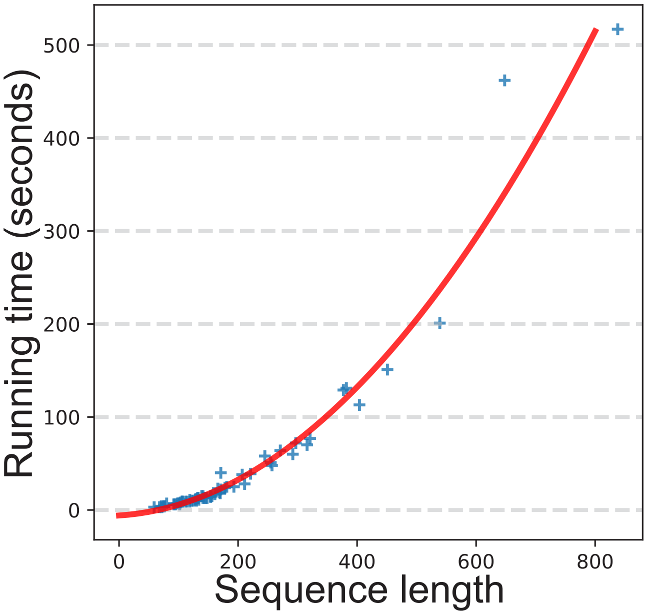


**Supplementary Figure 5. Running time of ProDESIGN-LE for protein sequence design**

The figure was plotted using the running time of ProDESIGN-LE for design sequences of the 68 CASP14 proteins.

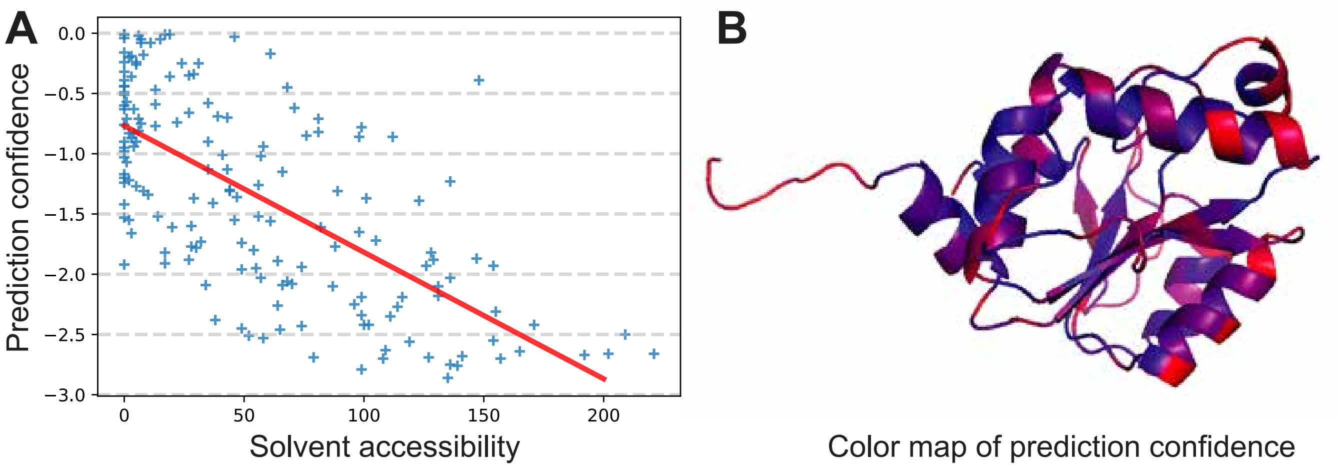


**Supplementary Figure 6. The relationship between the accuracy of the prediction of amino acid type and solvent accessibility of the target residue**

**A,** Using CASP14 domain T1045s2-D1 (166 a.a.) as an example, we plot the accuracy of the prediction of amino acid type and solvent accessibility of the target residue (Pearson correlation coefficient: $-0.68$). **B,** The color map of the prediction confidences on the 166 residues of T1045s2-D1. Red: low confidence, Blue: high confidence

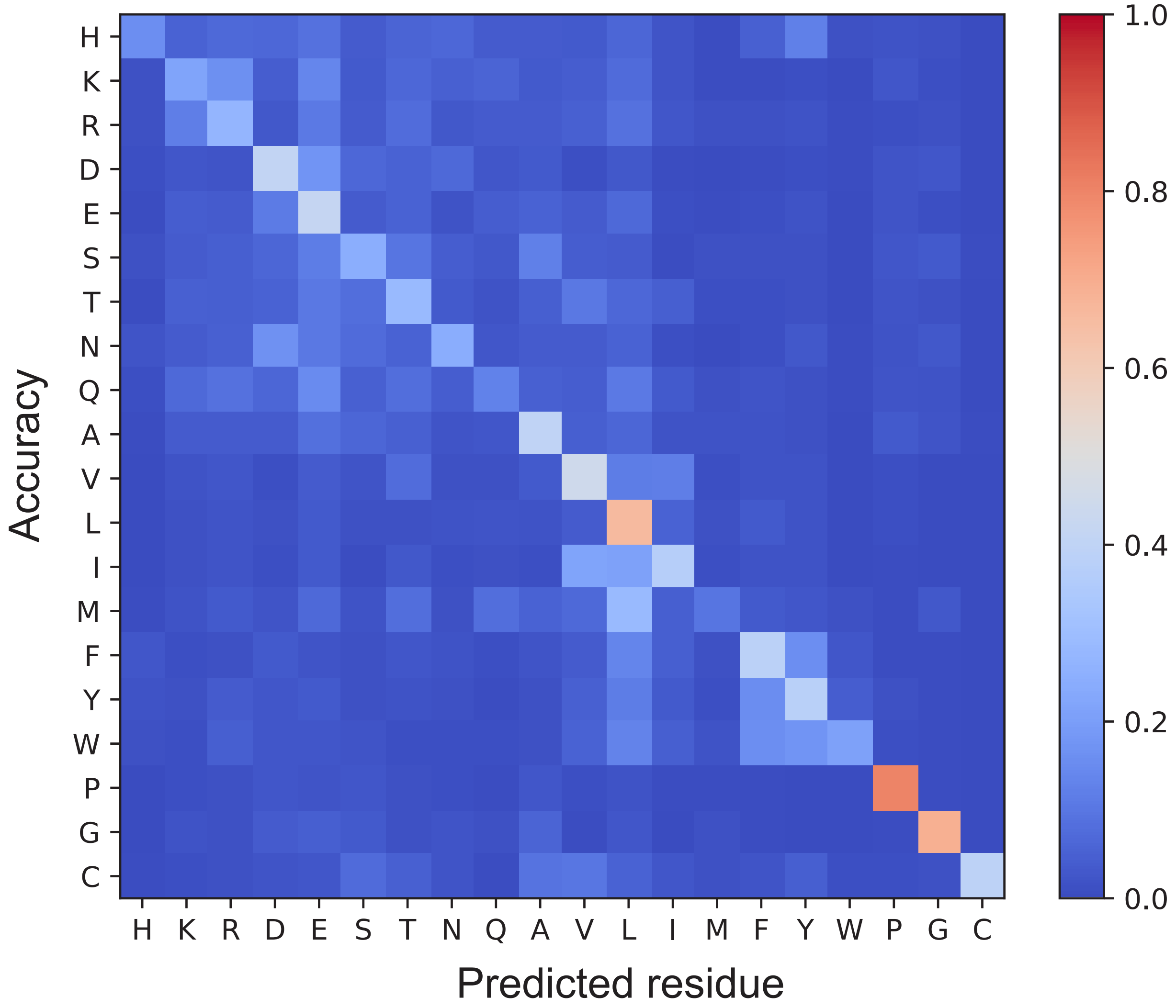


**Supplementary Figure 7. Confusion matrix of the transformer.**

The confusion matrix shows the ground-truth residue type and the predicted distribution over all possible 20 residue.

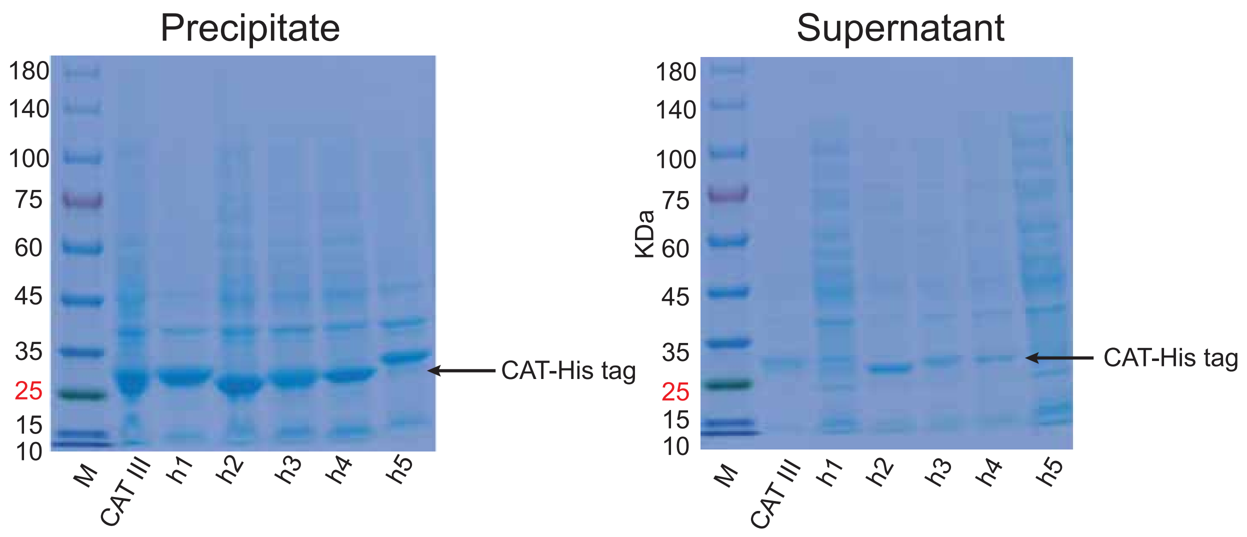


**Supplementary Figure 8. SDS-PAGE analysis of the five designed CAT proteins crude extracts**

**A,** SDS–PAGE analysis of the five designed CAT proteins precipitates induced at $16\,^{\circ}\mathrm{C}$.
**B,** SDS–PAGE analysis of the five designed CAT proteins supernatant induced at $16\,^{\circ}\mathrm{C}$. M: marker; CAT III: The natural protein (PDB: 6x7q). h1-h5: The five designed proteins. Extracted proteins were separated in $4$ to $12\,\%$ SDS-PAGE and stained with coomassie blue. Protein samples of $0.1\,\mathrm{OD}$ each were loaded on the gel. The molecular weight of the target protein is $28.3\,\mathrm{kDa}$.

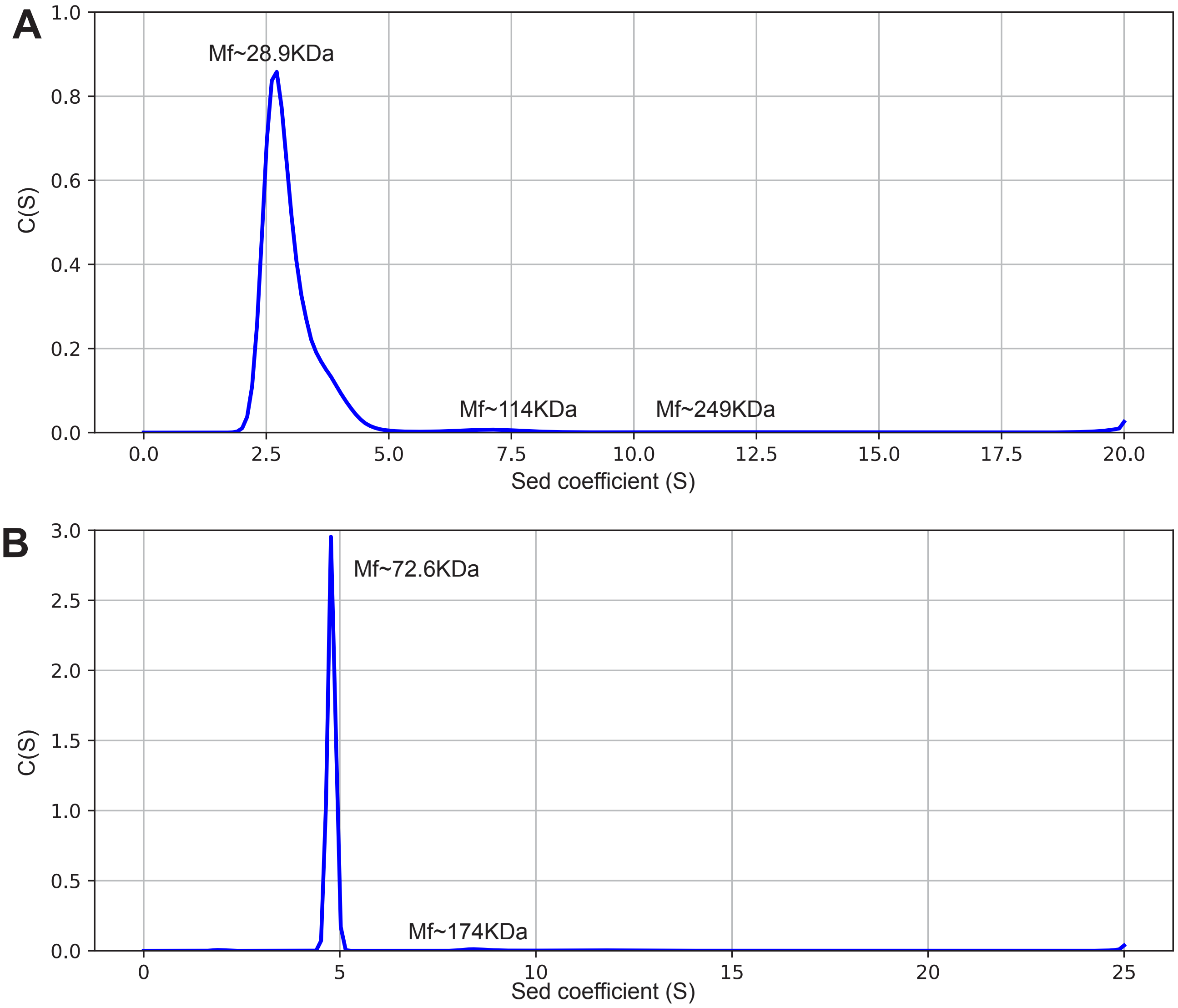


**Supplementary Figure 9. Sedimentation velocity analysis(SVA) of the designed CAT-h2 protein and the control CAT III protein**

A and B, according to Sedimentation velocity assay methods, obtained the subsidence coefficient (S) of the designed protein of CAT-h2(A) and the control protein of CAT III(B). **A,** The molecular weight (MW) of CAT-H2 protein $(28.9\,\mathrm{kDa}])$ is consistent with the MW of CAT-h2 monomer protein $(28.3\,\mathrm{kDa}])$ according to SVA. **B,** MW of control protein of CAT III $(72.6\,\mathrm{kDa}])$ is consistent with the MW of control trimer protein $(74.7\,\mathrm{kDa}])$ according to SVA.

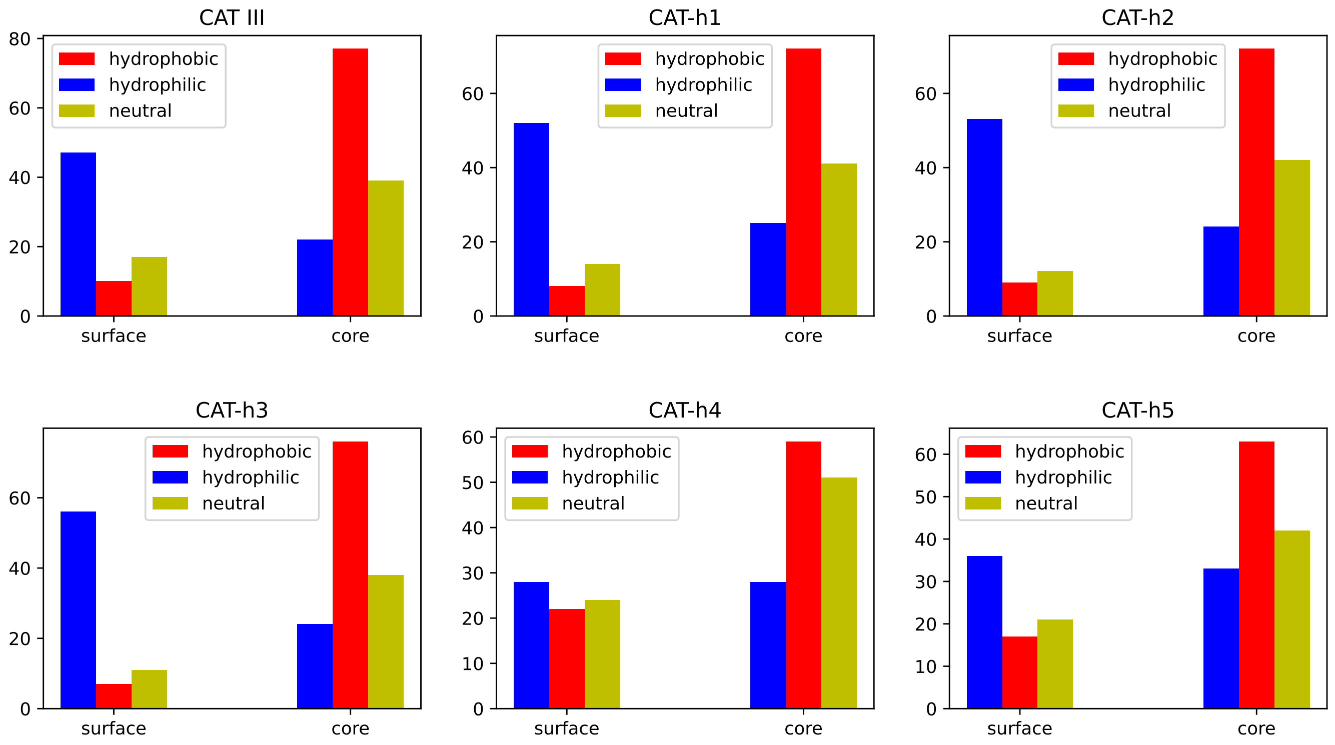


**Supplementary Figure 10. The ratios of hydrophobic, hydrophilic and neutral residues in the core or on the surface of the designed proteins for CAT III**
